# IDENTIFICATION OF CANDIDATE GENES AND PATHWAYS LINKED TO THE TEMPERAMENT TRAIT IN SHEEP

**DOI:** 10.1101/2023.10.12.562033

**Authors:** Estefanía Romaniuk, Brenda Vera, Pablo Peraza, Gabriel Ciappesoni, Juan Pablo Damián, Elize van Lier

## Abstract

Temperament can be defined as the emotional variability among animals of the same species in response to the same stimulus, thus being able to group animals by reactivity as nervous, intermediate or calm. The main objective for this study was to identify genomic regions with the temperament phenotype measured by the Isolation Box Test (IBT) by single-step genome wide association studies (ssGWAS). The database used consisted of 4,317 animals with temperament records, and 1,697 genotyped animals with 38,268 effective SNP after quality control. We identified three genomic regions that explained the greatest percentage of the genetic variance, resulting in 25 SNP associated with candidate genes on chromosomes 6, 10 and 21. A total of nine candidate genes are reported for the temperament trait, which are: *PYGM*, *SYVN1*, *CAPN1*, *FADS1*, *SYT7*, *GRID2*, *GPRIN3*, *EEF1A1* and *FRY*, linked to energetic activity of the organism, synaptic transmission, meat tenderness and calcium associated activities. This is the first study to identify these genetic variants associated with temperament in sheep, leading the opportinuty to potential applications of these molecular markers in future behavioral research.

## Introduction

Temperament is a tool to understand animal behavior and can be defined as the emotional variability among animals of the same species in response to the same stimulus (Blache and Bickell, 2010). Animals perceive environmental stimuli, such as novelty, uncertainty, challenge or change, and may react in distinct ways (Clarke and Boinski, 1995). Temperament refers to the individuals’ consistent behavioral styles or tendencies and can be represented in opposites like shy or bold, sociable or aggressive, restless or quiet (Southerland et al., 2012). Different physiological and behavioral responses have been used as indicators to classify animals according to their temperament (Veissier et al. 2009), and can be described as calm, nervous, or neutral. Temperament can influence productive traits such as ovulatory rate (Bickell et al. 2010; Van Lier et al. 2017), meat quality (Wulf et al. 2002), animal weight gain (Voisinet et al. 1997), milk and colostrum quality (Sart et al. 2004; Hawken et al. 2012), favoring animal welfare (Del Campo, 2011) and providing safety to the operator associated with a less reactive animals or of a calmer temperament (Haskell et al. 2014).

Variability in animal temperament may be influenced by polymorphism of genes that are expressed in the brain, or along the sympathic-adreno-medullary axis (SAM) of the autonomic nervous system and hypothalamus-pituitary-adrenal axis (HPA) (Qiu et al. 2016, 2017). The Single Nucleotide Polymorphism (SNP) is the most common type of polymorphism where a single base is changed for another (e.g. GGTACC/GGTGCC) at a frequency greater than 1 % in a population (Brookes, 1999). Since 1990, the University of Western Australia (UWA) has selected Merino sheep establishing lines of calm and nervous sheep (Qiu et al. 2017). In these temperament flock, eigth SNP were found to be distributed differently between calm and nervous sheep (Ding et al. 2020), associated with the serotonin, oxitocin and dopamine systems as well as the stress response axis according to their temperament (Qiu et al. 2016; Ding et al. 2020). In a recent review, Alvarenga et al. (2021) repored 148 genomic regions associated with 22 behavioral traits (Poissant et al., 2013; Hazard et al., 2014; Pant et al., 2016; Qiu et al., 2016) and 15 candidate genes in sheep.

Genome-wide association studies (GWAS) are used to identify genomic regions where candidate genes associated with a phenotypic trait are located. With the inclusion of genomics in selection programs (Aguilar et al. 2010; Misztal et al. 2014), a very powerful tool called Single Step Genomic Wide Association Study (ssGWAS) has been developed to identify SNP for the phenotype of interest. The ssGWAS method uses genomic breeding values (GEBV) based on the combined pedigree, genomic and phenotypic information and its conversion to SNP marker effect, adjusting for population structure at the same time (Legarra et al.. 2009; Misztal et al., 2014). The advantage of ssGWAS methods is that it integrates all pedigree, genomic and phenotypic information from genotyped as well as non-genotyped individuals in a one-step procedure, which allows the use of any model and all relationships simultaneously (Legarra et al., 2009; Lourenco et al., 2020). Through computational tools, the SNP effects, genetic variance and the p-value of SNP associated with temperament trait can be calculated (Wang et al. 2012). Therefore, we proposed to use this methodology to explore the temperament trait in Merino sheep in Uruguay.

The working hypothesis of this study was that there are SNP associated with the temperament trait in Australian Merino sheep. The main objective was to identify the SNP associated with the temperament phenotype that explained the greatest genetic variance and to subsequently do gene-set analyses to identify candidate genes, functional gene-sets and gene signaling pathways implicated in temperament of sheep.

## Results

### Records of temperament with the IBT

The frequency of agitation scores was concentrated to the left (skewed distribution), at low score values (Fig. 1). The mean agitation for the total population evaluated was 42, with a median of 37. The minimum score was 1 and the maximum was 190. Scores above the mean tended to spread out.

**Fig. 1.**
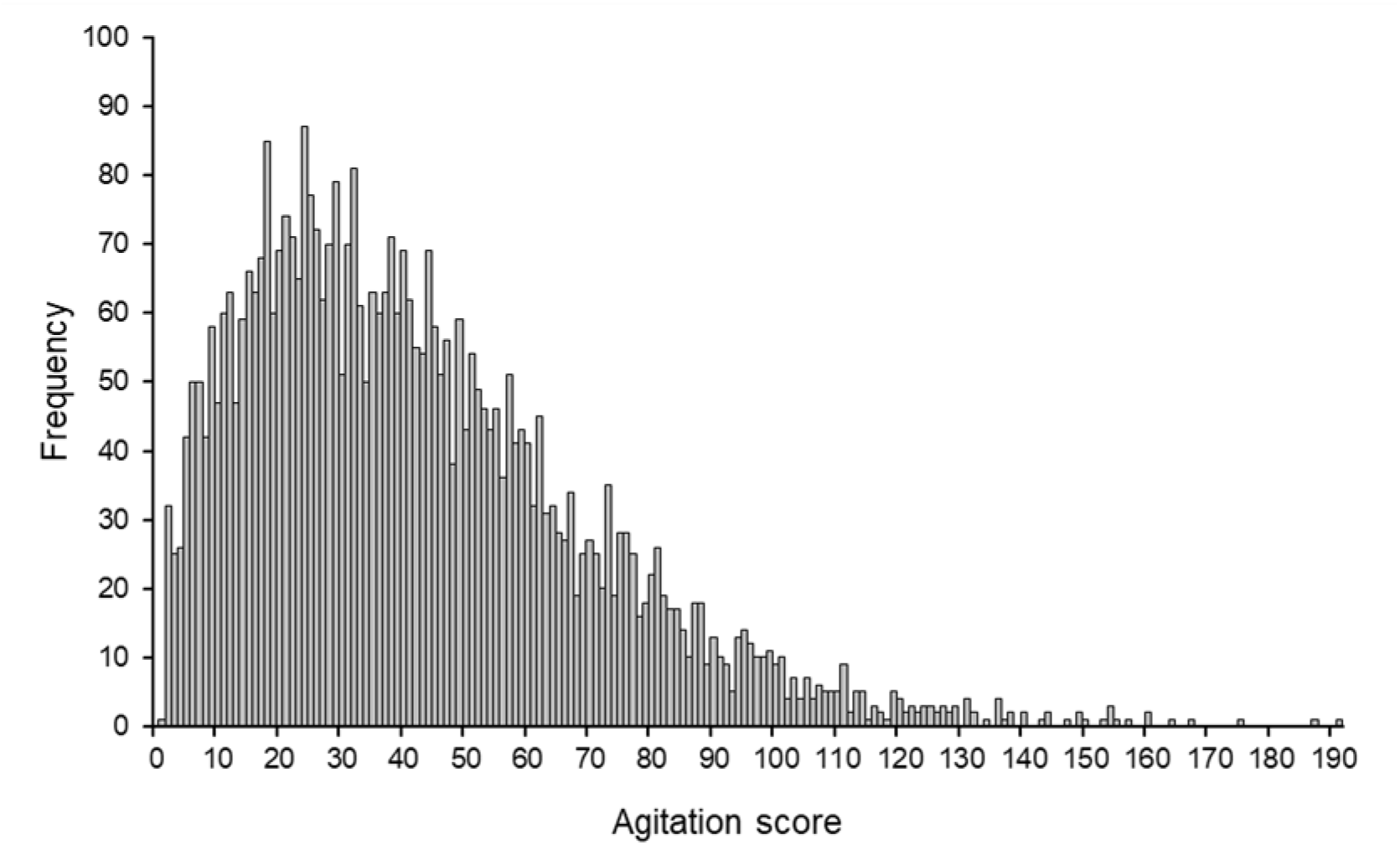
Frequency of agitation scores from the isolation box test for 4,317 Australian Merino lambs in response to social isolation for 30 seconds. The lower agitation scores represent less reactivity associated with a calmer temperament, and higher agitation scores represent more reactivity associated with nervous temperament.

### Estimation of genetic parameters

The genetic variance was 111.8 ± 22.8 (SEM) and the residual variance was 472.6 ± 20.80 (SEM). The heredability (h^2^) of the temperament trait was 0.19 ± 0.038 (SEM).

### Single Step Genome Wide Association Studies (ssGWAS)

The genetic variance explained by windows of 0.5 mbp of adjacent SNP, identified genomic regions of interest associated with chromosomes 6, 10 and 21 (var ≥ 0.5) (Fig. 2).

**Fig. 2.**
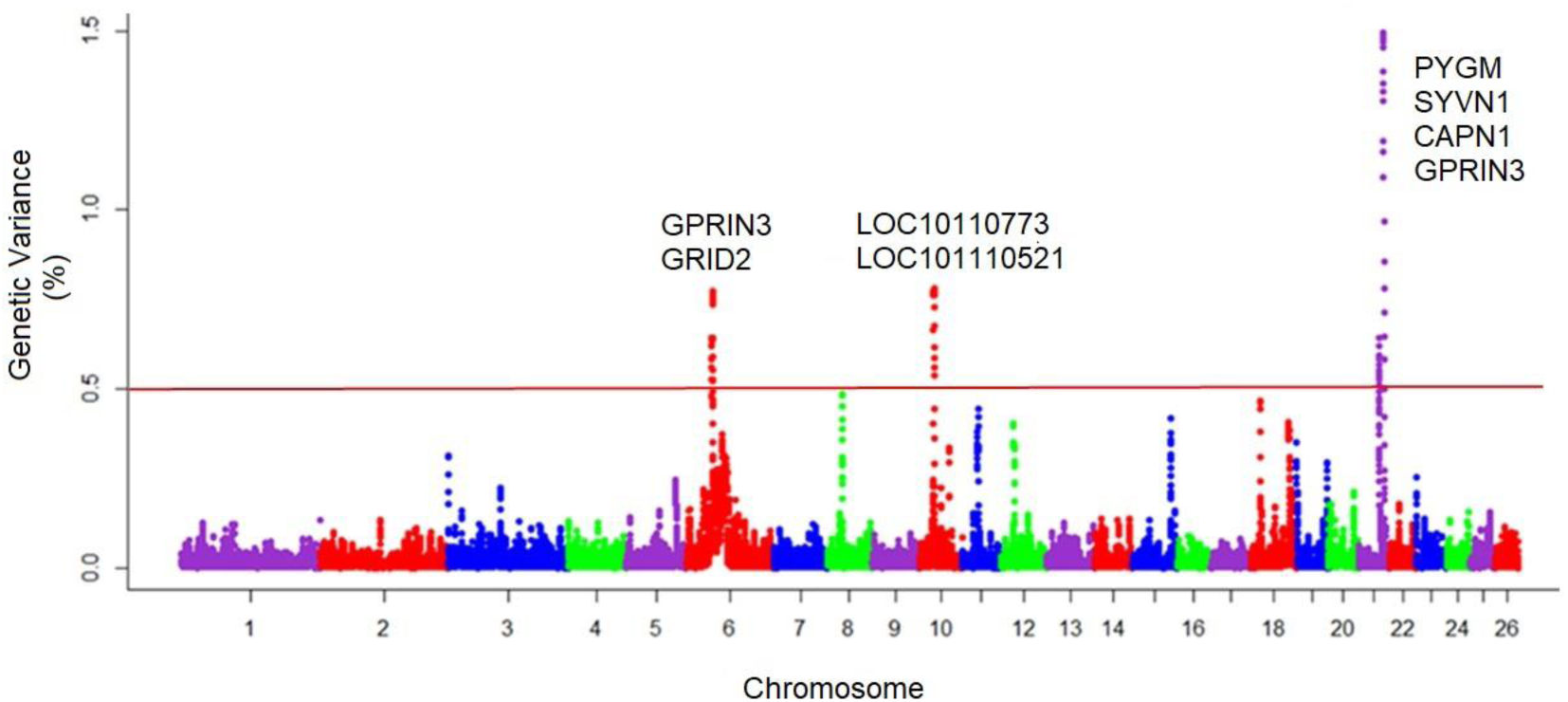
Manhattan plot of 38,268 effective SNP for 1,697 Australian Merino genotyped animals belonging to the National Genetic Evaluation, included in the database of the SULAR. Chromosome number and statistical significance threshold according to the % genetic variance explained by windows variance of 0.5 mbp. The solid horizontal line is the threshold, % var ≥ 0.5 %. The nine candidate genes involved in the temperament trait on chromosomes 6, 10 and 21 were inserted with their respective codes within each chromosome.

The information on the most relevant SNP found in these regions showed that there were nine candidate genes for the temperament trait (Table 1), which were: ***PYGM*** (21:42,295,599-42,307,126), ***SYVN1*** (21:42,650,559-42,655,527), ***CAPN1*** (21:42,712,976-42,740,799), ***LOC101110773*** (10:29,275,771-29,457,586), ***LOC101110521*** (10:28,986,741-29,188,660), ***GRID2*** (6:30,768,380-31,534,647), ***FADS1*** (21:39,652,537-39,665,108), ***SYT7*** (21:39,390,965-39,426,334) and the ***GPRIN3*** gene (6:35,511,293-35,513,635). The genes and annotation of the regions with more than 0.5 % of the genetic variance on chromosomes 6, 10 and 21 are shown in Table 2. On chromosome 21, the variant rs402505013 (21:42,295,749) determined the greatest genetic variance (1.46 %) and five candidate genes were detected. On chromosome 6 and 10, two candidate genes each were detected.

**Tab. 1.** SNP detected within the windows explaining more than 0.5 % of the genetic variance, sorted in descending order. Candidate genes associated with these SNP are reported in the next table.

| rs code | pos (bp) | CHR | % Var | Variant type | Candidate gene |
| --- | --- | --- | --- | --- | --- |
| rs402505013 | 42295749 | 21 | 1.4691 | Splice acceptor var | *PYGM* |
| rs419347404 | 42601748 | 21 | 1.4519 | Intron variant | - |
| rs161627521 | 42654067 | 21 | 1.3288 | Intron variant | *SYVN1* |
| rs413708295 | 42714381 | 21 | 1.1617 | Intron variant | *CAPN1* |
| rs421553713 | 42714613 | 21 | 1.0906 | Intron variant | *CAPN1* |
| rs161627624 | 42715850 | 21 | 0.8537 | Synonymous variant | *CAPN1* |
| rs408317317 | 29353089 | 10 | 0.7803 | Intron variant | *LOC101110773* |
| rs428995675 | 29072930 | 10 | 0.7738 | Intron variant | *LOC101110521* |
| rs407693533 | 35491698 | 6 | 0.7722 | Intron variant | - |
| rs422603241 | 31453177 | 6 | 0.7688 | Intron variant | *GRID2* |
| rs422288687 | 29304176 | 10 | 0.7678 | Intron variant | *LOC101110773* |
| rs427220269 | 29054709 | 10 | 0.7670 | Intron variant | *LOC101110521* |
| rs421383362 | 29188403 | 10 | 0.7636 | Intron variant | *LOC101110521* |
| rs409829992 | 29162222 | 10 | 0.7624 | Intron variant | *LOC101110521* |
| rs399382510 | 29202499 | 10 | 0.7619 | Intergenic variant | - |
| rs411759303 | 35511497 | 6 | 0.7430 | Missense variant | *GPRIN3* |
| rs400430030 | 29421760 | 10 | 0.6745 | Intron variant | *LOC101110773* |
| rs419116702 | 29030595 | 10 | 0.6646 | Intron variant | *LOC101110521* |
| rs398879843 | 35191867 | 6 | 0.6423 | Intergenic variant | - |
| rs161618576 | 39638962 | 21 | 0.6420 | Downstream gene variant | - |
| rs424142667 | 35511899 | 6 | 0.6415 | Missense variant | *GPRIN3* |
| rs410592527 | 35184703 | 6 | 0.6228 | Intergenic variant | - |
| rs408222545 | 39642723 | 21 | 0.6204 | Upstream gene variant | - |
| rs419203432 | 29415140 | 10 | 0.6173 | Intron variant | *LOC101110773* |
| rs161618641 | 39646260 | 21 | 0.5915 | Upstream gene variant | - |
| rs416558978 | 35254368 | 6 | 0.5873 | Intergenic variant | - |
| rs417392501 | 39651749 | 21 | 0.5832 | Downstream gene variant | - |
| rs404318469 | 42719476 | 21 | 0.5826 | Intron variant | *CAPN1* |
| rs403363266 | 39653383 | 21 | 0.5657 | 3 prime UTR variant | *FADS1* |
| rs419203432 | 29415140 | 10 | 0.5595 | Intron variant | *LOC101110773* |
| rs403382566 | 35512106 | 6 | 0.5508 | Missense variant | *GPRIN3* |
| rs421709693 | 39432569 | 21 | 0.5434 | Intergenic variant | *SYT7* |
| rs427110197 | 39654860 | 21 | 0.5384 | Intron variant | *FADS1* |
| rs398157763 | 29455959 | 10 | 0.5363 | 3 prime UTR variant | *LOC101110773* |
| rs399480023 | 31217615 | 6 | 0.5249 | Intron variant | *GRID2* |
| rs399060511 | 35275766 | 6 | 0.5247 | Intergenic variant | - |
| rs406335698 | 35276244 | 6 | 0.5219 | Intergenic variant | - |

rs = SNP reference; bp = base pairs, position of the SNP in the reference genome (versión oar\_v3.1); CHR = chromosome; % Var = percentage of the genetic variance; (-) = no candidate gene was detected for the SNP position

**Tab. 2.**
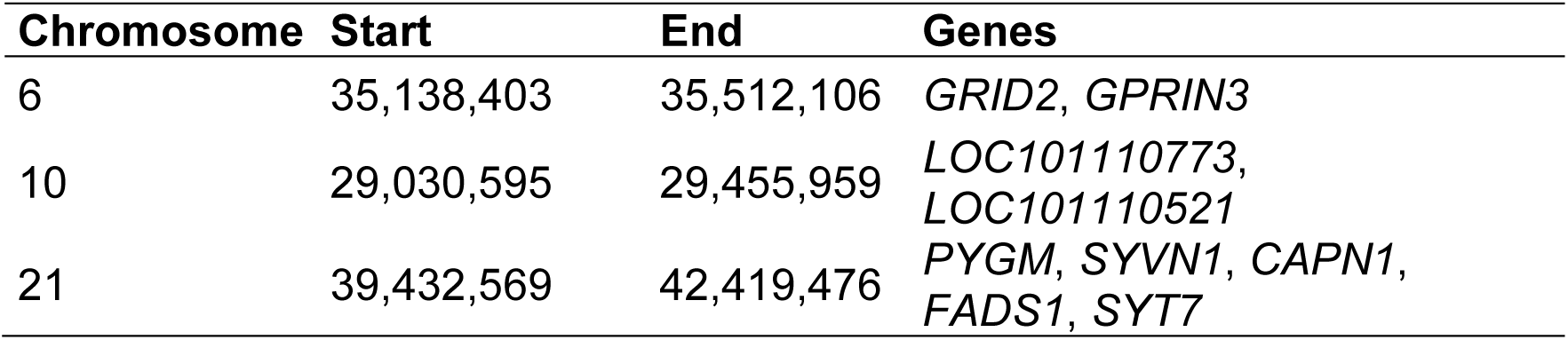
Genes located in the regions of the genome defined by windows of 0.5 mbp and that explain a genetic variance ≥ 0.5%, chromosome and its position in the sheep reference genome.

### Enrichment analysis

For the enrichment analysis 2,185 SNP were considered (5 % SNP greatest effect). This SNP defined a set of 900 genes in the sheep reference genome. The enriched p-value of each biological term indicated the importance of the term with respect to the set of analyzed genes (Table 3, Fig. 3). Several significant metabolic pathways (p-value ≤ 0.05) were identified and grouped into four categories: signaling, metabolism, steroidogenesis, and others (Table 3). The gene ontology of functional enrichment is shown in Figure 3. There were several terms linked to the regulation of ATP, calcium, cell activity, locomotion and behavior, lipid and carbohydrate metabolism, modulation of synaptic transmission, GTPase activity among others.

**Fig. 3.**
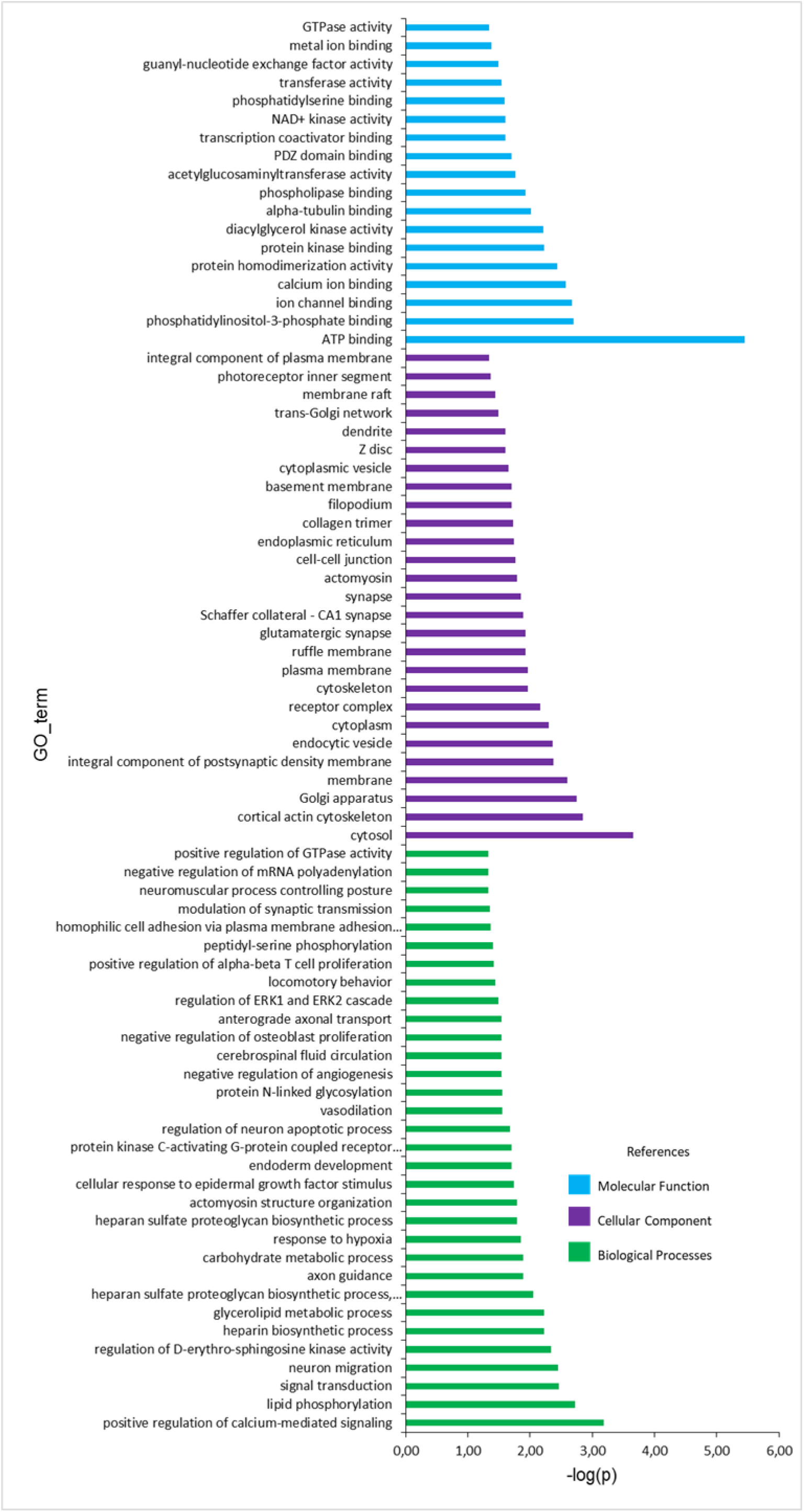
Gene ontology categories represented by Molecular Functions (light blue color), Cellular Component (purple color) and Biological Processes (green color), for the set of positional candidate genes with the 5 % of SNP with the greatest effect in function del –log of the enriched p-value.

**Tab. 3.**
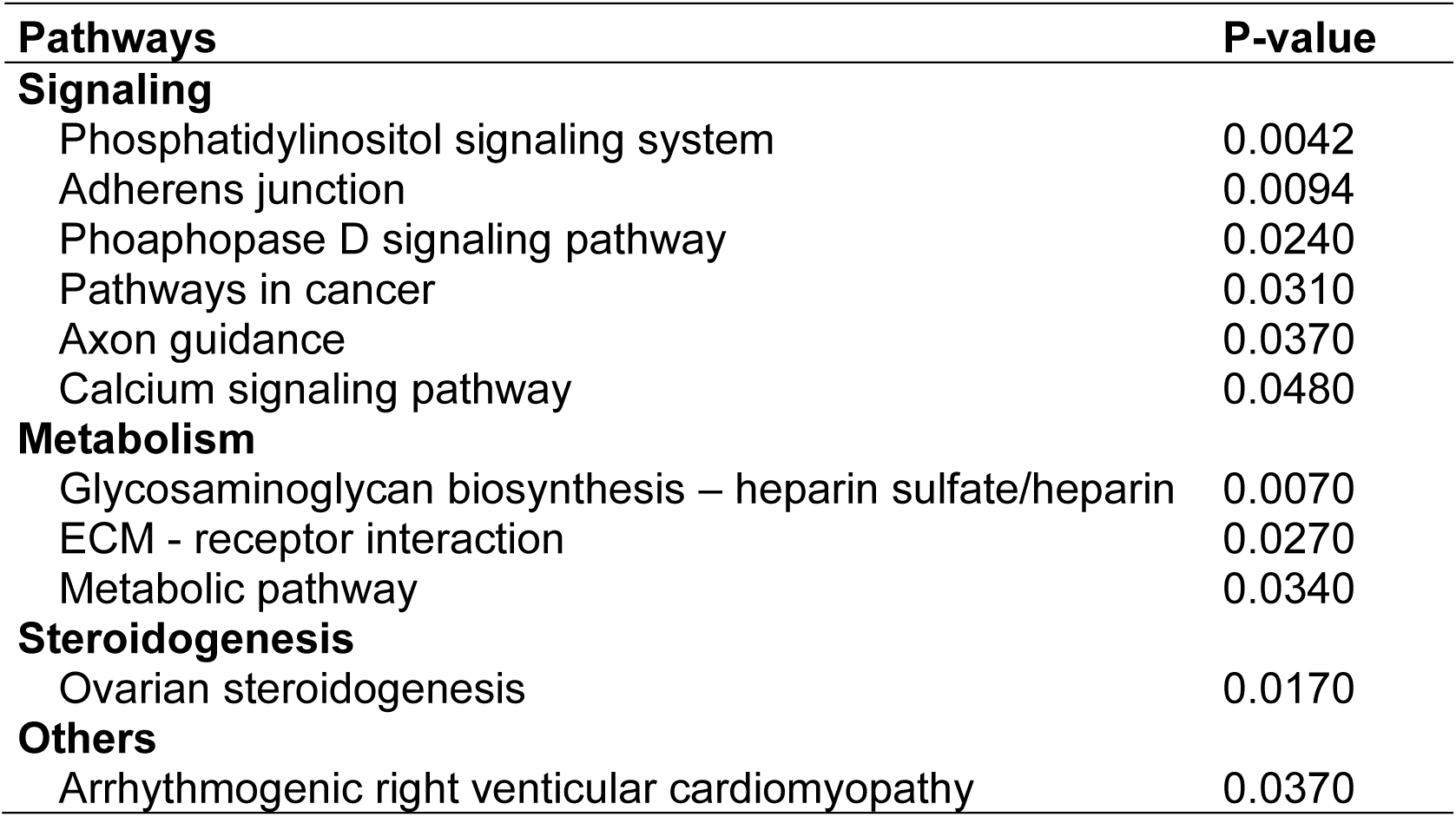
Detected pathways from the list of captured genes with the top 5 % SNP effect and their classification into 4 classes associated with the p-value of enrichment.

## Discussion

The hypothesis that there are SNP associated with the temperament trait in Australian Merino sheep was confirmed. The genetic variance of the SNP showed that there are several regions associated with greater variability, suggesting that temperament is not solely governed by the effect of one major gene. Instead, it appears to be a complex multigenic trait, influenced by multiple genes distributed across the genome, as suggested by Hazard et al. (2014). As part of the results of this study, nine positional candidate genes and gene signaling pathways underlying temperament in the Australian Merino sheep were detected, several of which have not been previously reported. These findings are valuable and relevant due to the following: 1) we worked with a population of animals that were not previously selected for temperament, 2) we utilized a molecular panel and information that included data reported by groups of Australian, European, and American researchers (GGP_50k_ovi, https://www.neogen.com/categories/genotyping-arrays/ggp-ovine-50k/), and 3) the size of the sample of genotyped animals and of the genealogy was appropriate for the methodology used. Therefore, the source of information to do the association study was reliable, robust and considered the structure of the population by including the genotyping and genealogy matrices.

### Heritability (h^2^)

The estimated heritability of temperament was 0.19 ± 0.038, which is a moderate h^2^ (Cardellino and Rovira, 1989). The h^2^ value shows that it would be feasible to include the temperament trait in a selection program. The estimated value (0.19) is consistent with the h^2^ for adult ewes (0.20, Plush et al., 2011) and is similar to the h^2^ for Corriedale lambs (0.18, Zambra et al., 2015). In another sheep study considering a subpopulation of the phenotypic data used in the present study, the h^2^ was 0.31 ± 0.06 (two generations, n = 2,952 animals) (Zambra et al., 2015). The difference in the estimated h^2^ value may be due to an increase in the number of measured individuals and some of the new animals are children of those that had been measured before. In the study by Zambra et al. (2015), there were no measurements of parents/children with data. Aditionally, the inclusion of genomic information from the animals, which translates into higher precision. On the other hand, in an Australian report that included more breeds of sheep, the h^2^ was higher: Australian Merino 0.38; Poll Dorset 0.41; White Suffolks 0.29; Poll Merino 0.41 (Blache and Ferguson, 2005a). The h^2^ for other productive species is similar to the estimated value (cattle: 0.22 – 0.26, Le Neindre et al., 1995 and Burrow et al., 1988; horses: 0.23, Oki et al., 2007). Despite the fact that temperament is not economically valued, several studies confirm its incidence in productive characters (Voisinet et al., 1997; Wulf et al., 2002; Sart et al., 2004; Bickell et al., 2010; Van Lier et al., 2017).

### Chromosomes and candidate genes

In chromosome 21, one SNP defined the greatest genetic variance and is within the *PYMG* gene (21:42,295,599-42,307,126) encodding the enzyme muscle glycogen phosphorylase, an allosteric enzyme that plays a central role in maintaining cellular and body glucose homeostasis (Tan et al., 1997). In a study about protein phosphorylation levels with different meat tenderness post-mortem, show that it is the main protein involved in the regulation of energy metabolism (Chen et al., 2015) and reported a positive correlation between the phosphorylation level of glycogen phosphorylase and the rate of glycolysis (Laville et al., 2009; Lametsch and Karlsson, 2012; Chen et al., 2015). The activity of this enzyme responds to stress stimuli and catalyzes the decomposition of glycogen to glucose 1-phosphate, which is converted to glucose 6-phosphate, the substrate for glycolysis (Lucia et al., 2008). In stressful situations such as social isolation, stress hormones like adrenaline, stimulate glycogen phosphorylase to quickly releasing glucose from muscle glycogen. The deficiency of this enzyme has been reported in humans and sheep as the cause of symptoms of intolerance to exercise, since less glucose would be available for muscle contraction (Tan et al., 1997; Lucia et al., 2008; Howell et al., 2014). A mutation in the *PYGM* gene has been reported in a flock of Merino sheep from Murdoch University Veterinary School farm in Western Australia (Tan et al., 1997; Walker, 2006). The mutation is an acceptor splice site in intron 19 of *PYGM* and determined an absence of glycogen phosphorylase activity in muscle fibres, double recessive animals by genotyping exhibit similar clinical effect to those seen in humans (Howell et al., 2014). One of the consequence of having less energy available for the organism negatively affects the stress response, where the main objective of the physiological changes is to increase the availability of energy in response to the stressor (Sapolsky et al., 2000; Damián et al., 2015). On the other hand, glycogen phosphorylase is one of the most important enzymes involved in the tenderness of meat and may affect the quality of it (Bai et al., 2020a). Therefore, genetic variations could not only affect what is related to stress or welfare, but the stress conditions linked to the end of the productive stage will determine the characteristics of the final product, in this case the meat (Bai et al., 2020a, 2020b).

Variants rs413708295 (21:42,714,381), rs421553713 (21:42,714,613) and rs161627624 (21:42,715,850), identified Calpain gene, subunit 1(*CAPN1*) (21:42,712,976-42,740,799). This gene encodes the μ-calpain 1, a proteolytic enzime with activity on myofibrillar proteins (Speck et al., 1993; Page et al., 2002; Geesink et al., 2006). This gene is associated with proteolytic proteins directly linked to the tenderness of meat and are essential for postmortem proteolysis in the process of transforming muscle into meat (Speck et al. 1993). Two SNP located in exons 9 and 14 in a Piedmontese-Angus bull resulted in an amino acid substitution in the *CAPN1* protein. The change from cytokine to guanine in exon 9 caused the substitution of alanine for glycine at position 316 of the protein sequence (A^316^/G^316^), while the change of guanine to alanine determined the substitution of valine for isoleucine at position 530 (V^530^/I^530^) (Page et al., 2002). Analysis of genotypes revealed that (G^136^/I^530^) alleles produced less tender meat compared to genotypes with (A^136^/V^530^) alleles (Page et al., 2002). In Santa Ines sheep, four SNP (rs401662939, rs407017992, rs418468486 and rs420860201) had suggestive additive effects on eight meat quality traits: rib eye area, water holding, internal carcass length, conformation carcass score, leg length, leg yield, meat lightness and yellowness (Meira et al., 2020). In another study with the same sheep, variants in the *CAPN1* gene were associated with weaning weight (rs417258958), rib eye area (rs403953588 and rs430307080), fat thickness (rs408790217), body depth (rs420860201), and heights at croup and withers (rs408790217) (Machado et al., 2020), all of these traits were recorded *in vivo*. That variants in these genes were also associated with physicochemical meat traits such as pH, color, tenderness and water-holding capacity (Meira et al., 2020). The differential gene expression of *CAPN1* among animals of different temperaments might be involved a lower meat quality for the more reactive animals, which needs further research for confirmation.

The rs161627521 (21:42,654,067) variant is within the Synoviolin gene (*SYVN1*) (21:42,650,559-42,655,527). The *SYVN1* gene has been extensively studied in relation to body weight regulation and mitochondrial biogenesis. According to Fujita et al. (2015), *SYVN1* is an E3 ligase that negatively regulates the thermogenic coactivator peroxisome proliferator activated receptor coactivator (PGC)-1β. Also codes for the W5PTU3 protein (RING-type E3 ubiquitin transferase) linked to the ubiquitination and protein modification pathways (The UniProt Consortium, 2021). To our knowledge, this is the first study documenting variants in the *SYVN1* gene in sheep. More studies are necessary to underly the implications of this gene in sheep and their productive characteristics. Another gene identified by two markers rs403363266 (21:39,653,383) and rs427110197 (21:39,654,860), was the Fatty Acid Desaturase 1 gene (*FADS1*) (21:39,652,537-39,665,108), associated with the W5PWA9 protein in sheep (The UniProt Consortium, 2021). The *FADS1* gene is a member of the fatty acid dehydrogenase family and is related to all lipids (Laan et al., 2018). This gene is part of a set of genes significantly enriched in biosynthesis of the unsaturated fatty acid signaling pathway, fatty acid metabolism, adipocyte differentiation, and other processes related to fat storage (Liu et al., 2022). The last candidate gene associated with the variant rs421709693 (21:39,432,569) was Synaptotagmin 7 (*SYT7*) (21:39,390,965-39,426,334). The *SYT7* gene has been studied in relation to synaptic transmission (Jackman et al., 2016) and behavioral abnormalities (Chen et al., 2017). According to Chen et al. (2017), is involved in ensuring the efficiency of high-frequency transmission at central GABAergic synapses. The study found that genetic elimination of *SYT7* reduced the frequency of asynchronous release during high-frequency stimulus trains. However, the function of *SYT7* in synaptic transmission at GABAergic synapses was found to be less prominent compared to other synapses (Chen et al., 2017). In terms of temperament, there is limited research specifically linking the *SYT7* gene to this trait. However, a study by Shen et al. (2020) investigated the behavioral phenotype of mice with silenced genes, including *SYT7*, in the hippocampus. They observed manic-like and depressive-like behavioral fluctuations in a majority of *SYT7* knockout mice, which were analogous to the mood cycling symptoms of bipolar disorder. This suggests that *SYT7* may be a candidate risk factor for behavioral abnormalities (Shen et al., 2020). The molecular functions and biological processes linked to this gene are associated with calcium (e.g. regulation, dependent activation for the fusion of synaptic vesicles, exocytosis of neurotransmitters, repair of the plasmatic membrane, regulation of dopamine secretion, glucagon and insulin secretion, among others) (The UniProt Consortium, 2021). The main enzymes of proteolysis depend on calcium (Koohmaraie, 1992) and anything related to plasma calcium concentrations could be involved in meat quality. On the other hand, there are genes that have been associated with behavioral traits and that are also found within or near the genomic region of interest on chromosome 21. This is the case of the *CHRM1* gene (21:37.9-48.4 Mb), muscarinic cholinergic receptor, which would be linked to locomotion, cognition and the nervous system (Tanda et al., 2007). Therefore, the SNP of chromosome 21 found in the present study could be used as molecular markers for future studies focused on candidate genes associated with temperament or behavioral traits that affect reactivity and that are associated with meat quality.

The variants rs408317317 (10:29,353,089), rs422288687 (10:29,304,176), rs400430030 (10:29,421,760), rs419203432 (10:29,415,140) and rs398157763 (10: 29,455,959) identified the gene LOC101110773 (*EEF1A1*) (10:29,275,771-29,457,586) on chromosome 10, linked to molecular functions of energetic activity. Specifically, the encoded protein W5PD15 (elongation factor alpha 1) promotes GTP-dependent binding of aminoacyl-tRNA to the A-site of ribosomes during protein biosynthesis (The UniProt Consortium, 2021). Another gene detected on chromosome 10 was the LOC101110521 (10: 28,986,741-29,188,660) and the five associated markers were rs427220269 (10: 29054709), rs421383362 (10: 29,188,403), rs4098299992 (10: 29,162,222) and rs419116702 (10:29,030,595). This gene on the Ensembl platform is described as the *FRY* gene. A phenotypic study in Merino and Merino derived sheep reported this gene as one of the major signals in the genome linked to the phenotypic traits of wool (Megdiche et al., 2019). In this same study, the *EEF1A1* gene (LOC101110773) was also described as part of a set of genes of biological interest that contributes to elucidating the genetic basis of the Merino phenotype. Two SNP have been reported for temperament trait (Ding et al., 2020) in Merino sheep and the associated gene is 5 – hydroxytryptamina receptor 2A (HTR2A) in chromosome 10 (rs17196799, rs7193181) (Ding et al., 2020). Half of the SNP of chromosome 10 that contribute to the genetic variance of temperament were associated with molecular functions of energetic activity. This is consistent with aspects of reactivity, such as physiological response to stress, where the animal recognizes the threat to homeostasis and biological responses are activated to restore normal homeostasis function and welfare (Moberg, 2000).

On chromosme 6, the region of interest with more than 0.5 % of the genetic variance was between base pairs 35,138,403 and 35,512,106. In this region, the *GPRIN* candidate gene of subfamily number 3 (*GPRIN3*) (6:35,511,293-35,513,635) associated with the variants rs111759303 (6:35,511,497), rs424142667 (6:35,511,899) and rs403382565 (6:35,511,899) was identified. This gene is highly expressed in the adrenal gland and codes for the W5NQE9 protein, but actually there is not a lot of information about it. However, in a study about neuronal excitability, morphology and striatal-dependent behaviors in the indirect pathway of the striatum, Karadurmus et al. (2019) founded that *GPRIN3* was involved as a mediator of dopamine receptor D2 and was assosiated with schizophrenia, parkinson’s disease, and drug addiction. I addition, how *GPRIN3* gene is expressed in the adrenal gland, it is a relevant finding given that the adrenal gland is involved in the stress response (SAM and HPA axes, Selye, 1939; Moberg, 2000; Sapolsky et al., 2000) and there are differences in the frequency of polymorphism according to the type of temperament (Qiu et al., 2016). The A/A genotype of the SNP628 type polymorphism of the *CYP17* gene specifically involved in cortisol production was more frequent in sheep sellected for nervous temperament, while the G/G genotype was in sheep sellected for calm temperament (Qiu et al., 2016).

The other candidate gene on chromosome 6 is *GRID2* (6:30,768,380-31,534,647) associated with two markers, rs399480023 (6:31,217,615) and rs422603241 (6:31,453,177). This gene codes for the W5QA32 protein (glutamate receptor), which is related to biological processes linked to the regulation of synaptic transmission (The UniProt Consortium 2021). Particulary, the glutamate receptor functions as an ion channel in the central nervous system and plays a very important role in transmission excitation (The UniProt Consortium 2021). Glutamate is an essential aminoacid, is the most abundant neurotransmitter in the brain, also involved in the regulation of behavioral, social, learning, memory, sensory and cognitive processes (Giménez et al., 2018). It is a neurotransmitter involved in reproduction due to its favorable action on hypothalamic neurons for the secretion of GnRH (Iremonger et al., 2010). According to Iremonger et al. (2010), Glutamato regulates sexual behavior through dopamine, due to its action on hypothalamic GnRH neurons. Recent research in sheep demonstrated that the inclusion of specific amino acids in the diet, such as arginine, glutamine, leucine and glycine, has beneficial effects on embryonic and fetal survival and growth (Wu, 2010; Saevre et al., 2011; Crane et al., 2016). On the other hand, in a study to identify QTL (Quantitative Trait Locus) for behavioral reactivity revealed a QTL that maps within the gene (Glutamate receptor metabotropic 7), associated with locomotion in response to social isolation (Hazard et al., 2014). In addition, this QTL is close to the gene encoding the oxytocin receptor (OXTR) which is associated with social (Sala et al., 2013; Damián et al., 2021) and maternal behaviors (Champagne et al., 2001; Feldman et al., 2012). An study support this revealing a SNP rs17664565 in chromosome 19 maps within the gene OXTR (Ding et al., 2020).

Therefore, the SNP detected on chromosome 6 identified two relevant genes, one that is expressed in the axis linked to the stress response (HHA axis, *GPRIN3* gene) and another that is expressed centrally (*GRID2* gene) associated to behavioral traits. Temperament is key in both scenarios, since greater reactivity negatively affects the functioning of the reproductive endocrine axis (Dobson and Smith, 2000; Damián et al., 2015).

### Functional enrichment analysis

Enrichment analysis revealed several functional pathways and biological processes associated with the temperament trait. The gene-set is associated with a set of annotation terms. If the genes share a similar set of those terms, they are most likely involved in similar biological mechanisms. The DAVID platform provides an algorithm that adopts Kappa statistics, which enables the quantitatively measurement of the degree of agreement of genes that share similar anotation terms. The Kappa result ranges from zero to one (higher value, higher agreement). In the calcium signaling pathway (p-val = 0.048) several genes were grouped, but the most important were: *HTR5A* gene, serotonin receptor; *DRD1* gene, dopamine receptor and *GRM1* gene, glutamate receptor 1. A more detailed exploration of calcium signaling was assosiated the following annotation terms: cardiac muscle contraction (kappa = 0.42); oxytocin signaling (kappa = 0.41); glutamate synapse (kappa = 0.35) and dopamine synapse (kappa = 0.31). The neurotransmitters dopamine and serotonin are involved in the stress response in a variety of species, through the activation of their different receptors (Reif and Lesch, 2003; Noblett and Coccaro, 2005). Oxytocin is an important regulator of social behaviors (Damián et al., 2021) such as maternal behavior (Meyer-Lindenberg et al., 2011) and recognition in humans (Kumsta et al., 2013), and polymorphism in the oxytocin receptor gene has been associated with temperament, reactivity to stressors, and aggressive behaviors (Rodríguez et al., 2009; Tost et al., 2010; Malik et al., 2012). Therefore, the gene-set revealed that the enriched calcium signaling pathway is closely related to temperament because involves many aspects of this trait and its physiological effects, as has been shown previously in reproductive aspects (Bickell et al., 2010; Van Lier et al., 2017).

Five genes were identified in the metabolic pathway of steroidogenesis in the ovary (p-val = 0.0170): *CYP2J* gene (cytochrome P450); follicle stimulating hormone receptor gene (*FSHR*); gene for the enzyme hydroxysteroid 17-beta dehydrogenase (*HSD17B2*) and two genes associated with prostaglandin synthesis (*PGFS*, *PTGS2*). These genes linked to reproduction reaffirm the association of temperament with reproduction and its effect on the reproductive endocrine axis: hypothalamic-pituitary-gonadal (HHG) (Dobson and Smith, 2000; Von Borell et al., 2007; Damián et al., 2015).

In the metabolic pathways, the *ACTN2* gene was found to be associated to arrhythmogenic cardiomyopathy of the right ventricular (p-val = 0.037). This gene codes for the protein actinin alpha 2, reported for humans, which is expressed in both skeletal muscles and the heart, and its function is the anchoring of thin myofibrillar actin filaments (Beggs et al., 1992; Ribeiro et al., 2014; Murphy and Young, 2015). A variant in the regulatory region of this gene leads to heart failure (Arvanitis et al., 2020). However, to our knowledge, this is the first report of its association with the temperament trait in sheep.

One of the enriched biological processes considered the term positive regulation of calcium-mediated signaling (GO:0050850, p-val = 0.001), where the FSH receptor gene and others (*CD24*, *CDH13*, *SYK*, *LOC101112639*) were observed. The term locomotor behavior (GO:0007626, p-value = 0.036) considers the specific movement of an animal from one place to another in response to external or internal stimuli and/or a combination of the internal state and external conditions of that animal. A subset of this GO_term is directly linked to the behavior term (GO:0007610), described as an animal’s responses to internal or external stimuli (actions or inactions), through a mechanism that involves the activity of the nervous system. Another GO_term regulation of ERK1 and ERK2 cascade (Extracellular Signal Regulated Kinase) (GO:0070372, p-val = 0.032) participates in various biological responses. This term is associated with several processes that modulate the frequency, rate, or extent of signal transduction mediated by the ERK1 and ERK2 cascade. This signaling cascade is a pathway formed by mitogen-activated protein kinases that regulate a wide variety of cellular processes, as diverse as proliferation, differentiation, survival, and stress (Herrero, 2012). The GO_term linked to carbohydrate metabolic processes (GO:0005975, p-val = 0.013) is also revealed by the association that presents the availability of energy to cope with a stressor, specifically the physiological responses to stress that include an increase in metabolic rate and energy, as well as an increase in lipid catabolism and protein degradation. These physiological changes in the animal help to maintain the inmediate availability of energy and oxygen (Sapolsky et al., 2000; Damián et al., 2015), and is the basis of the “fight or flight” response, the first autonomous response of the individual to face the stressor or to escape from it (Cannon, 1914).

Our results revealed SNP associated with the temperament trait in sheep. These SNP were not previously reported, which opens a window for exploration, not only linked to temperament but also to different metabolic pathways that could have an impact on production and welfare. The genetic variance of the SNP revealed that there are regions of the genome associated with greater variability and that temperament is not regulated by the effect of a major gene, but rather is a multigenic characteristic. Nine genes were detected in the genomic regions on the chromosome 6, 10 and 21 (genes: *GRID2, GPRIN3, LOC101110773, LOC101110521, PYGM, SYVN1, CAPN1, FADS1, SYT7*), linked to the energetic activity of the organism, synaptic transmission, meat tenderness and calcium associated activities. The identification of these genes, metabolic pathways and their respective functions should contribute to a better understanding of the genetic mechanisms that regulate the temperament trait in sheep. The functional enrichment analysis identified gene functions and complemented the results of the association study. The 5 % of the SNP with the greatest effect determined a set of 900 genes linked to various metabolic pathways, where the gene ontology analysis shows that there are several processes linked to the regulation of ATP, calcium, cell activity, locomotion and behavior, lipid and carbohydrate metabolism, modulation of synaptic transmission, among others. Several of these metabolic pathways are of interest for the temperament trait but need further exploration.

## Materials and methods

### Phenotypic and pedigree data

A total of 4,317 lambs were tested for temperament between one and three months post-weaning, including four progenies: 2010, 2011, 2018 and 2019. The records of temperament scores were included in the genetic database (SULAR, Uniform System of Survey and Records Storage, of the National Genetic Evaluation). Temperament was measured using the IBT (Blache and Ferguson 2005a, 2005b). The dimensions of the box were 1.50 m (L) x 1.50 m (H) x 0.75 m (W). Each lamb was gently pushed inside the box, held there for 30 seconds, and agitation was objectively measured by an agitation meter. The agitation meter registered the vibrations of the box induced by the lamb’s movements and high-pitched vocalizations. The higher the agitation score, the more nervous the sheep. The average age of the evaluated lambs was 160 ± 38 days at the time of the IBT test.The IBT was calibrated with an electronic unit to high and low agitation score. The intra-test coefficients of variation (CV %) of the IBT were 7.17 % and 7.93 % for the high and low setting, respectively. The inter-test CV % of the IBT were 9.37 % and 9.29 % for the high and low setting, respectively. There was no previous selection of animals and they had no previous experience with the IBT. All procedures were approved by the Honorary Commission for Animal Experimentation (Comisión Honoraria de Experimentación Animal, CHEA, protocols N° 021130-006307-11 and 020300-000653-18-ID, CHEA 701), Universidad de la República.

Genealogical information from 10,799 animals were provided by the Rural Association of Uruguay and the Merino Breeders Society. For the genotyped animals, it was checked that there were no parent-offspring incompatibilities based on Mendelian conflict counts using the SeekParentF90 software (Misztal et al., 2014). To verify the consistency of the information, a Pearson correlation was performed between the off-diagonal elements of the genomic and pedigree information matrices.

### Genotypic data and quality control

DNA was extracted from blood samples following the protocol described by Medrano et al. (1990). A total of 1,697 lanimals were genotyped for 43,705 SNP and the molecular information was obtained using the Geneseek Genotypong Profile panel (GeneSeek ® Genotyping Profile, GGP, Illumina, San Diego, California). Genomic data quality control (QC) was conducted using PREGSF90 (Misztal et al., 2014) included the exclusion of markers located in sexual chromosomes, monomorphic SNP, with minor allele frequencies (MAF) < 0.05% and call rates < 85%. Animals were removed from the analysis when call rates were < 90 %. A total of 38,268 effective SNP were retained for subsequent genomic analyses.

### Model and estimation of genetic parameters

The mixed model to perform a ssGWAS included fixed effects (genetic and environmental) and random effects. The fixed effects were contemporary group (year, flock, sex and management group), dam age (< 2 years, 2 to 3 years or > 3 years) and type of birth (single or twin). The age of the lamb at the time of measurement with the IBT was considered as a covariate.

The following model was used (Zambra et al., 2015):

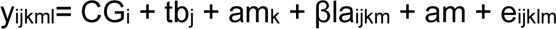

where: **y** is the temperament data of the lamb m; **CG**_i_ is the contemporary group (i: year-flock-sex-management group; 73 levels), **tb**_j_ the type of birth of lamb (j: single (1), twin (2)), **am**_k_ the age of dam subdivided into 3 classes (k_1_ = < 2 years ewe, k_2_ = 2 to 3 years ewe and k_3_ = > 3 years ewe), **β** is the regression coefficient of the age of the lamb on temperament, **la** is the covariate of lamb age (in days) at the time of measurement, **a** is the additive effect of the animal with distribution **am** ∼ N (0, σ^2^_A_ A) where σ^2^_A_ is the additive variance of temperament, A is an additive relationship matrix, and **e** is the random effect of the error with independent normal distribution between the observations with variance σ^2^e.

The variance components necessary for the model were estimated by airemlf90 program version 1.148 of BLUPF90.

The heredability was calculated as:

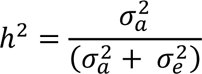

where 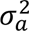 is the genetic variance and 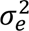 is the residual variance.

### Single Step Genome Wide Association Studies (ssGWAS)

The ssGWAS methodology allows genome-wide association studies to be performed by combining all available pedigree, phenotype, and genotype information in a single evaluation using the BLUPf90 family of programs (Misztal et al., 2014). The result of combining the genomic (SNP) and pedigree information is the [H] matrix, where the molecular information of genotyped animals is projected, through relatives, to individuals that are not genotyped (Misztal et al., 2014). Here, the classical pedigree relationship [A] matrix is replaced by [H] matrix. Thus, all the SNP are considered simultaneously together with all the phenotypes of the genotyped and non-genotyped animals (Aguilar et al., 2010; Wang et al., 2014). The [H] matrix is complex, it can be simplified to its inverse [H^-1^] (Aguilar et al., 2010):

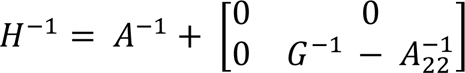

where G^-1^ is the inverse of genomic relationship matrix (it is for genotyped animals, proportion of alleles shared between animals); A^-1^ is the inverse of pedigree relationship matrix; and A_22_^-1^ is the inverse of the pedigree relationship matrix for genotyped animals. The [G] matrix was computed according to VanRaden (2008):

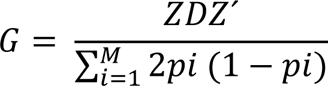

where Z is an incidence matrix adjusted for allele frequencies; D is a diagonal matrix of weights for SNP variances; Z’ is the transpose; M is the number of SNP; and p_i_ represents the minor allele frequence of each SNP.

The percentage of genetic variance (% var) explained by a region has been calculated as follows (Wang et al., 2014):

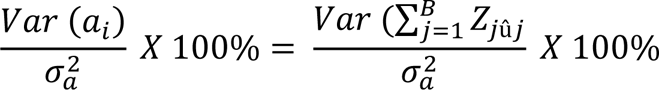

where a_i_ is genetic value of the i-th region; B is the total number of adjacent SNP within 0.5 Mb region; 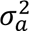 is the total genetic variance; Z_j_ is vector of gene content of the j-th SNP for all individuals, and 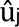 is marker effect of the j-th SNP within the i-th region.

The SNP effects were estimated according to Stranden and Garrik (2009):

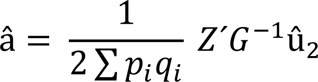

where Ź is the transpose of Z matrix; G is the genomic matrix for genotyped animal; 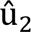 is the genomic breeding value for genotyped animals and p_i_q_i_ are the allele frequencies.

### Identification of positional candidate genes

The threshold non-overlapping windows that explain more than 0.5 % genetic variance (Brunes et al., 2020; Sigdel et al., 2021) were reported and were considered as candidate regions to verify overlapping genes. The SNP that explain a greater genetic variance in the window were reported. Manhattan plot based on the proportion of additive genetic variance explained by the windows were generated using the R.Script output from POSTGSF90 (Misztal et al., 2014).

### Gene enrichment analysis

On the other hand and only for this approach the 5 % of SNP with the greatest effect were selected (Stranden and Garrik, 2009; Sigdel et al., 2021; Jara et al., 2022). An SNP was assigned to a gene if it was located inside a gene or within a genomic distance of 5 kb upstream or downstream from a gene (Brunes et al., 2021; Sigdel et al., 2021). To assign SNP to genes, computational procedures were used with an ad hoc script in the R programmer (version 4.1.0) (R Core Team, 2021), gene locations were obtain using BiomaRt package (version 2.48.3) (Durinck et al., 2005, 2009) and the Ovis_aries_v3.1 ovine reference genome data set (https://www.ensembl.org; Cunningham et al., 2022). Therefore, a gene was associated with temperament if it contained at least one SNP whose effect was in the top 5 % of the distribution. To assign the genes to functional categories, DAVID (Database for Annotation, Visualization and Integrated Discovery, latest available version 6.8, Sherman et al., 2022) was used. The objective of this analysis was to know in which metabolic pathways the genes would be involved and quantify it through the enriched p-value. The available Gene Ontology (GO) terms for molecular function (MF), cellular component (CC) and biological process (BP) were included in the analysis, and Kyoto Encyclopedia of Genes and Genomes (KEGG) databases were used to reveal the functional implication of the detected genes. Finally, the significance of the terms enriched was a threshold p-value ≤ 0.05, as implemented in the DAVID platform.

## CRediT authorship contribution statement

**Romaniuk Estefanía**: Investigation, Data Curation, Formal analysis, Writing – Original draft, Writing – Review & Editing, **Vera Brenda**: Validation, Formal analysis, Writing – Original draft, Writing – Review & Editing, **Peraza Pablo**: Resources, Methodology, Formal analysis, Writing – Review & Editing, **Ciappesoni Gabriel**: Conceptualization, Methodology, Investigation, Writing – Review & Editing, Project administration, Funding acquisition, **Damián Juan Pablo**: Conceptualization, Writing – Review & Editing, Project administration, Funding acquisition, **Van Lier Elize**: Conceptualization, Investigation, Writing – Original draft, Writing – Review & Editing, Supervision, Funding acquisition. Conflict of interest

The authors declare no competing interests.

## Acknowledgement

This work is supported by grants from the Comisión Sectorial de Investigación Científica (CSIC, Universidad de la República), which approved the project CSIC_I+D_2018_287; the SMARTER project funded by the European Union Horizon 2020 program (agreement no. 772787); the projects INIA CL 40 (resistance to parasites) and CL 38 (RUMIAR); and to the Comité Académico de Posgrados for the scholarship master’s desree 2020 (POS_NAC_2019_1_157979), Universidad de la República.

## References

Aguilar, I., Misztal, I., Johnson, D.L., Legarra, A., Tsuruta, S., Lawlor, T.J., 2010. Hot topic: A unified approach to utilize phenotypic, full pedigree, and genomic information for genetic evaluation of Holstein final score. J Dairy Sci. 93(2), 743–752. doi: 10.3168/jds.2009-2730.

Aguilar, I., Misztal, I., Tsuruta, S., Legarra, A., Wang, H., Legarra, A., 2020. PREGSF90 – POSTGSF90: Computational Tools for the Implementation of Single-step Genomic Selection and Genome-wide Association with Ungenotyped Individuals in BLUPF90 Programs. 10th World Congress on Genetics Applied to Livestock Production (WCGALP), Aug 2014, Vancouver, Canadá. American Society of Animal Sciense. Proceedings. https://hal.inrae.fr/hal-02743809

Alvarenga, A.B., Oliveira, H.R., Chen, S.Y., Miller, S.P., Marchant-Forde, J.N., Grigoletto, L., Brito, L.F., 2021. A Systematic Review of Genomic Regions and Candidate Genes Underlying Behavioral Traits in Farmed Mammals and Their Link with Human Disorders. Animals 2021, 11, 715. 10.3390/ani11030715

Arvanitis, M., Tampakakis, E., Zhang, Y., 2020. Genome-wide association and multi-omic analyses reveal ACTN2 as a gene linked to heart failure. Nat Commun. 11: 1122. 10.1038/s41467-020-14843-7

Bai, Y., Li, X., Zhang, D., Chen, L., Hou, C., Zheng, X., Ren, C., Ijaz, M., 2020a. Role of phosphorylation on characteristics of glycogen phosphorylase in lamb with different glycolytic rates post-mortem. Meat Science, 164. 10.1016/j.meatsci.2020.108096

Bai, Y., Li, X., Zhang, D., Chen, L., Hou, C., Zheng, X., Ren, C., 2020b. Effects of phosphorylation on the activity of glycogen phosphorylase in mutton during incubation at 4 °C in vitro. Food Chemistry, 313. 10.1016/j.foodchem.2020.126162

Beggs, A.H., Byers, T.J., Knoll, J.H., Boyce, F.M., Bruns, G.A., Kunkel, L.M., 1992. Cloning and characterization of two human skeletal muscle alpha-actinin genes located on chromosomes 1 and 11. JBC. 267(13), 9281–9288.

Bickell, S., Durmic, Z., Blache, D., Vercoe, P.E., Martin, G.B., 2010. Rethinking the management of health and reproduction in small ruminants. Updates on Ruminant Production and Medicine. 26th World Buiatrics Congress. Proceedings. 14–17 November 2010, Santiago, Chile.

Blache D, Ferguson D. 2005a. Genetic Estimates for Temperament traits in sheep Breeds. Reproduction, 364.

Blache, D., Ferguson, D., 2005a. Genetic Estimates for Temperament traits in sheep Breeds. Rep. 364. 19 p.

Blache, D., Ferguson, D., 2005b. Increasing sheep meat production efficiency and animal welfare by selection for temperament meat. MLA. 364: 1–22

Blache, D., Bickell, S.L., 2010. Temperament and reproductive biology: emotional reactivity and reproduction in sheep. Rev Bras Zoo. (39): 401-408.

Brookes, A.J., 1999. The essence of SNPs. Gene. 234: 177–186. doi: 10.1016/S0378-1119(99)00219-X.

Brunes, L.C., Baldi, F., Lopes, F.B., Lôbo, R.B., Espigolan, R., Costa, M.F., Stafuzza, N., Magnabosco, C.U., 2020. Weighted single-step genome-wide association study and pathway analyses for feed efficiency traits in Nellore cattle. J Anim Bree Genet. 138(1), 23–44. doi:10.1111/jbg.12496.

Burrow, H.M., 2001. Variances and covariances between productive and adaptive traits and temperament in a composite breed of tropical beef cattle. Livestock Production Science 70, 213–233.

Cannon, W.B., 1914. The emergency function of the adrenal medulla in pain and the major emotions. Am J Physiol. 33(2), 356–372.

Cardellino R, Rovira J. 1989. Mejoramiento genético animal. Montevideo: Hemisferio Sur. 253.

Cardellino, R., Rovira, J., 1989. Heredabilidad. In Mejoramiento genético animal (ed. R Cardelino and J Rovira), pp. 111–151. Hemisferio Sur, Montevideo, Uruguay

Champagne, F., Diorio, J., Sharma, S., Meaney, M.J.., 2001. Naturally occurring variations in maternal behavior in the rat are associated with differences in estrogen-inducible central oxytocin receptors. Proc Natl Acad Sci U S A 2001, 98(22):12736–12741.

Chen, L., Li, X., Ni, N., Liu, Y., Chen, L., Wang, Z., Zhang, D., 2016. Phosphorylation of myofibrillar proteins in post-mortem ovine muscle with different tenderness. Journal of the Science of Food and Agriculture, 96(5), 1474–1483.

Chen, C., Satterfield, R., Young, S., Jonas, P., 2017. Triple Function of Synaptotagmin 7 Ensures Efficiency of High-Frequency Transmission at Central GABAergic Synapses. 21(8), 2082-2089. doi:10.1016/j.celrep.2017.10.122

Clarke, A.S., Boinski, S., 1995. Temperament in nonhuman primates are not genotyped. Am J Primatol. 37(2), 103–25. doi:10.1002/aip.1350370205

Crane, A.R., Redden, R.R., Van Emon, M.L., Neville, T.L., Reynolds, L.P., Caton, J.S., Schauer, C.S., 2016. Impacts of supplemental arginine on the reproductive performance of fall lambing ewes. J Anim Sci. 94: 3540–3549. ISSN: 1525-3163. doi: 10.2527/jas.2016-0379

Cunningham, F., Allen, J.E., Allen, J., Alvarez-Jarreta, J., Ridwan, M., Armean, I., Irimoloye, O.A., Azov, A.G., Barnes, I., Bennett, R., Berry, A., Bhai, J., Bignell, A., Billis, K., Boddu, S., Brooks, L., Charkhchi, M., Cummins, C., Da Rin Fioretto, L., Davidson, C., Dodiya, K., Donaldson, S., El Houdaigui, S., El Naboulsi, T., Reham, F., Garcia Giron, C., Genez, T., Gonzalez, J., Guijarro-Clarke, C., Gymer, A., Hardy, M., Hollis, Z., Hourlier, T., Hunt, T., Juettemann, T., Kaikala, V., Kay, M., Lavidas, I., Le, T., Lemos, D., Marugán, J.C., Mohanan, S., Mushtaq, A., Naven, M., Ogeh, D., Parker, A., Parton, A., Perry, M., Piližota, I., Prosovetskaia, I., Sakthivel, M.P., Abdul Salam, A.I., Schmitt, B.M., Schuilenburg, H., Sheppard, D., Pérez-Silva, J.G., Stark, W., Steed, E., Sutinen, K., Sukumaran, R., Sumathipala, D., Suner, M.M., Szpak, M., Thormann, A., Tricomi, F.F., Urbina-Gómez, D., Veidenberg, A., Walsh, T.A., Walts, B., Willhoft, N., Winterbottom, A., Wass, E., Chakiachvili, M., Flint, B., Frankish, A., Giorgetti, S., Haggerty, L., Hunt, S.E., IIsley, G.R., Loveland, J.E., Martin, F.J., Moore, B., Mudge, J.M., Muffato, M., Perry, E., Ruffier, M., Tate, J., Thybert, D., Trevanion, S.J., Dyer, S., Harrison, P.W., Howe, K.L., Yates, A.D., Zerbino, D.R., Flicek, P., Ensembl., 2022. Nucleic Acids Research. 50, Issue D1, 7 January 2022, Pages D988–D995. doi:10.1093/nar/gkab1049

Clarke, A.S., Boinski, S., 1995. Temperament in nonhuman primates. Am J Primatol. 37(2), 103–125. doi:10.1002/ajp.1350370205

Damián, J.P., Bausero, M., Bielli, A., 2015. Acute stress, hypothalamic-hypophyseal-gonadal axis and testicular function-a review. Ann Anim Sci. 15(1): 31–50. doi:10.2478/aoas-2014-0084

Damián, J.P., De Soto, L., Espindola, D., Gil, J., Van Lier, E., 2021. Intranasal oxytocin affects the stress response to social isolation in sheep. Physiol & Behav. 230: 8. doi:10.1016/j.physbeh.2020.113282.

Del Campo, M., 2011. El bienestar animal en nuestros sistemas de poducción, enfoque de la investigación a nivel nacinoal. ln: INIA Tacuarembó. Unidad Experimental Glencoe. Día de campo, setiembre 2011, Paysandú, Uruguay. Propuestas tecnológicas para el incremento de la productividad, la valorización y el ingreso económico para sistemas ganaderos de basalto. Tacuarembó (Uruguay): INIA, 2011. p. 59-60.

Ding, L., Maloney, S., Wang, M., Rodger, J., Chen, L., Blanche, D., 2020. Association between temperament related traits and single nucleotide polymorphisms in the serotonin and oxytocin systems in Merino sheep. Genes, Brain and Behavior. Pages 1–32. 10.21203/rs.3.rs-34713/v1

Dobson, H., Smith, R.F., 2000. What is stress, and how does it affect reproduction? Anim Rep Sci. 60–61, 743–752

Durinck, S., Moreau, Y., Kasprzyk, A., Davis, S., De Moor, B., Brazma, A., Huber, W., 2005. BioMart and Bioconductor: a powerful link between biological databases and microarray data analysis. J Leukocyte Biol. 21(16): 3439–3440. 10.1093/bioinformatics/bti525

Durinck, S., Spellman, P., Birney, E., Huber, W., 2009. Mapping identifiers for the integration of genomic datasets with the R/Bioconductor package biomaRt. Nat Protoc. 4: 1184–1191.

Feldman, R., Zagoory-Sharon, O., Weisman, O., Schneiderman, I., Gordon I., Maoz, R., Shalev, I., Ebstein, R.P., 2012. Sensitive parenting is associated with plasma oxytocin and polymorphisms in the OXTR and CD38 Genes. Biol Psychiatry 2012, 72(3):175–181. doi:10.1186/1471-2164-15-778.

Fujita, H., Yagishita, N., Aratani, S., Saito-Fujita, T., Morota, S., Yamano, Y., Hansson, M., Inazu, M., Kokuba, H., Sudo, K., Sato, E., Kawahara, K., Nakajima, F., Hasegawa, D., Higuchi, I., Sato, T., Natsumi, A., Usui, C., Nishioka, K., Nakatani, Y., Maruyama, I., Usui, M., Hara, N., Uchino, H., Elmer, E., Nishioka, K., Nakajima, T., 2020. The E3 ligase synoviolin controls body weight and mitochondrial biogenesis through negative regulation of PGC-1β. The EMBO Journal. 34 (8), 1042–1055. doi:10.15252/embj.201489897.

Geesink, G., Kuchay, S., Chishti, A., Koohmaraie, M., 2006. Micro-calpain is essential for postmortem proteolysis of muscle proteins. Journal of Animal Science 84, 2834–2840.

Giménez, C., Zafra, F., Aragón, C., 2018. Fisiopatología de los transportadores de glutamato y de glicina: nuevas dianas terapéuticas. Rev Neurol. 67(12): 491–504. doi: 10.33588/rn.6712.2018067

Haskell, M.J., Simm, G., Turner, S.P., 2014. Genetic selection for temperament traits in dairy and beef cattle. Front Genet. 5: 1–18. doi:10.3389/fgene.2014.00368

Hazard, D., Moreno C., Foulquié D., Delval E., Francois D., Bouix, J., Sallé., Boissy, A., 2014. Identification of QTLs for behavioral reactivity to social separation and humans in sheep using the OvineSNP50 BeadChip. BMC Genomic. 15(1). Pages 1-18

Hawken, P., Williman, M., Milton, J., Kelly, R., Nowak, R., Blache, D., 2012. Nutritional supplementation during the last week of gestation increased the volume and reduced the viscosity of colostrum produced by twin bearing ewes selected for nervous temperament. Small Rum Res. 105(1–3): 308–314. doi: 10.1016/j.smallrumres.2012.01.011

Herrero, A., 2012. Especificidad espacio-temporal de las señales Ras-ERK en la determinación de respuestas biológicas. Tesis Doc. Bioquímica. Cantabria, España. Instituto de Biomedicina y Biotecnología de Cantabria.

Howell, J., Walker, K., Creed, K., Dunton, E., Davies, L., Quinlivan, R., Karpati, G., 2014. Phosphorylase re-expression, increase in the force of contraction and decreased fatigue following notexin-induced muscle damage and regeneration in the ovine model of McArdle disease. Neuromuscular Disorders. 24(2): 167–177. 10.1016/j.nmd.2013.10.003

Iremonger, K.J., Constaintin, S., Liu, X., Herbison, A.E., 2010. Glutamate regulation of GnRH neuron excitability. Brain Res. 1364: 35–43. doi:10.1016/j.brainres.2010.08.071

Jackman, S.L., Turecek, J., Belinsky, J.E., Regehr, W.G., 2016. The calcium sensor synaptotagmin 7 is required for synaptic facilitation. Nature. 529:88–91. doi: 10.1038/nature16507

Jara, E., Peñagaricano, F., Armstrong, E., Ciappesoni, G., Iriarte, A., Navajas, A., 2022. Revealing the genetic basis of eyelid pigmentation in Hereford cattle. J Anim Sci. 100(5). doi:10.1093/jas/skac110

Karadurmus, D., Rial, D., De Backer, J.F., Communi, D., Exaerde, A., Schiffmann, S., 2019. GPRIN3 Controls Neuronal Excitability, Morphology, and Striatal-Dependent Behaviors in the Indirect Pathway of the Striatum. Journal of Neuroscience. 39(38), 7513–7528. doi:10.1523/JNEUROSCI.2454-18.2019.

Koohmaraie, M., 1992. Effect of pH, temperature, and inhibitors of autolysis and catalytic activity of bovine skeletal muscle m-calpain. J. Anim. Sci. 70:3071

Kumsta, R., Heinrichs, M., 2013. Oxytocin, stress and social behavior: neurogenetics of the human oxytocin system. Curr Opin Neurobiol. 23, 11–6. 10.1016/j.conb.2012.09.004

Laan, S., Harshfiel, E., Hemerich, D., Stacey, D., Wood, A., Asselbergs, F., 2018. From lipid locus to drug target through human genomics. Cardiovascular Research. 114(9), 1258–1270. 10.1093/cvr/cvy120.

Lametsch, R., Karlsson, A., 2012. Electrical stimulation affects metabolic enzyme phosphorylation, protease activation and meat tenderization in beef. Journal of Animal Science, 90(5), 1638–1649

Laville, E., Sayd, T., Morzel, M., Blinet, S., Chambon, C., Lepetit, J., Hocquette, J., 2009. Proteome changes during meat aging in tough and tender beef suggest the importance of apoptosis and protein solubility for beef aging and tenderization. Journal of Agricultural and Food Chemistry, 57, 10755–10764.

Le Neindre, P., Trillat, G., Sapa, J., Ménissier, F., Bonnet, J.N., Chupin, M., 1995. Individual differences in docility in Limousin cattle. Journal of Animal Science 73, 2249–2253

Legarra, A., Aguilar, I., Misztal, I., 2009. A relationship matrix including full pedigree and genomic information. J Dairy Sci. 92, 4656–4663.

Liu, T., Feng, H., Youduf, S., Xie, L., Miao, X., 2022. Differential regulation of mRNAs and IncRNAs related to lipid metabolism in Doulang and Small Tail Han sheep. Scientific reports. pp 1–12. 10.1038/s41598-022-15318-z4.

Lucia, A., Nogales-Gadea, G., Pérez, M., Martín, M., Andreu, A., Arenas, J., 2008. McArdle disease: what do neurologists need to know? Nat Rev Neurol 4, 568–577. 10.1038/ncpneuro0913.

Lourenco, D., Legarra, A., Tsuruta, S., Masuda, Y., Aguilar, I., and Misztal, I., 2020. Single-step genomic evaluations from theory to practice: using snp chips and sequence data in blupf90. Genes, 11(7), 1–32. 10.3390/genes11070790.

Machado, A. l., Meira, A.N., Muniz, E.N., Azevedo, H., Coutinho, I., Mourão, G., Pedrosa, V., Pinto, I., 2020. Single loci and haplotypes in CAPN1 and CAST genes are associated with growth, biometrics, and in vivo carcass traits in Santa Inês sheep. Annals of Animal Science 10.2478/aoas-2020-0007.

Malik, A.I., Zai, C.C., Abu, Z., Nowrouzi, B., Beitchman, J.H., 2012. The role of oxytocin and oxytocin receptor gene variants in childhood-onset aggression. Genes Brain Behav. 11, 545–51. doi:10.1111/j.1601-183X.2012.00776.x.

Medrano, J.F., Aasen, E., Sharrow, L., 1990. DNA extraction from nucleated red blood cells. Biotechniques 8(1), 43.

Megdiche, S., Mastrangelo, S., Hamouda, M., Lenstra, J., Ciani, E., 2019. Merino and Merino-derived sheep breeds: a further look at genome-wide selection signatures for wool traits. Front Genet. 10. doi:10.3389/fgene.2019.01025.

Meyer-Lindenberg, A., Domes, G., Kirsch, P., Heinrichs, M., 2011. Oxytocin and vasopressin in the human brain: social neuropeptides for translational medicine. Nat Rev Neurosci. 12, 524–38. doi:10.1038/nrn3044.

Meira, A., Jucá, A., de Souza, T., Coutinho, L., Mourão, G., Azevedo, H., Muniz, E., Pedrosa, V., Batista, L., 2020. Post-mortem carcass traits are associated with μ-calpain and calpastatin variants in santa inês sheep. Animal Science Papers and Reports. Institute of Genetics and Animal Biotechnology of the Polish Academy of Sciences, Jastrzębiec, Poland. 38(4), pp. 369–380

Misztal, I., Aguilar, I., Legarra, A., Tsuruta, S., Lourenco, D.A., Fragomeni, B., Zhang, X., Muir, W.M., Cheng, H.H., Wing, T., Hawken, R.R., Zumbach, B., Fernando, R., 2020. GWAS using ssGBLUP I. 10th World Congress of Genetics Applied to Livestock Production GWAS, 5, 1–6. doi:10.1109/CGames.2013.6632596.

Misztal, I., Tsuruta, S., Lourenco, D., Masuda, Y., Aguilar, I., Legarra, A., Vitezica, Z., 2014. Manual for BLUPF90 family of programs.

Moberg, G.P., 2000. Biological response to stress: implications for animal welfare. In: Moberg G.P and Mench J.A (editors). The biology of animal stress: basic principles and implications for animal welfare. CABI Publishing, pp. 1–22. Wallingford, Oxon, UK.

Murphy, A.C., Young, P.W., 2015. The actinin family of actin cross-linking proteins - a genetic perspective. Cell Biosci. 5(49). doi:10.1186/s13578-015-0029-7.

Noblett, K.L., Coccaro, E.F., 2005. Molecular genetics of personality. Curr Psychiatry Rep. 7(1): 73–80. doi:10.1007/s11920-005-0028-1.

Oki, H., Kusunose, R., Nakaoka, H., Nishiura, A., Miyake, T., Sasaki, Y., 2007. Estimation of heritability and genetic correlation for behavioural responses by Gibbs sampling in the thoroughbred racehorse. Journal of Animal Breeding and Genetics 124, 185–191

Page, B.T., Casas, E., Heaton, M.P., Cullen, N.G., Hyndman, D.L., Morris, C.A., Crawford, A.M., Wheeler, T.L., Koohmaraie, M., Keele, J.W., Smith, T.P., 2002. Evaluation of single-nucleotide polymorphisms in CAPN1 for association with meat tenderness in cattle. J Anim Sci. 80(12), 3077–3085. doi: 10.2527/2002.80123077x.

Pant, S.D., You, Q., Schenkel, L.C., Voort, G.V., Schenkel, F.S., Wilton, J., Cain, L., Karrow, N.A., 2016. A genome-wide association study to identify chromosomal regions influencing ovine cortisol response. Livest. Sci. 187, 40–47.

Plush, K., Hebart, M., Brien, F., Hynd, P., 2011. The genetics of temperament in Merino sheep and relationships with lamb survival. Applied Animal Behaviour Science 134, 130–135.

Poissant, J., Réale, D., Martin, J.G.A., Festa-Bianchet, M., Coltman, D.W.A., 2013. Quantitative trait locus analysis of personality in wild bighorn sheep. Ecol. Evol. 3, 474–481.

Qiu, X., Ledger, J., Zheng, C., Martin, G.B., Blache, D., 2016. Associations between temperament and gene polymorphisms in the brain dopaminergic system and the adrenal gland of sheep. Physiol & Behav. 153, 19–27. doi:10.1016/j.physbeh.2015.10.022.

Qiu, X., Martin, G.B., Blache, D., 2017. Gene polymorphisms associated with temperament. J Neurogenetics. 31(1–2), 1–16. doi:10.1080/01677063.2017.1324857.

R Core Team, 2019. R: A language and environment for statistical computing. R Foundation for Statistical Computing, Vienna, Austria. URL https://www.R-project.org/.

Reif, A., Lesch, K.P., 2003. Toward a molecular architecture of personality. Behav Brain Res. 139(1-2), 1–20. doi: 10.1016/S0166-4328(02)00267-X.

Ribeiro, E.A., Pinotsis, N., Ghisleni, A., Salmazo, A., Konarev, P.V., Kostan, J., Sjöblom, B., Schreiner, C., Polyansky, A.A., Gkougkoulia, E.A., Holt, M.R., Aachmann, F.L., Zagrović, B., Bordignon, E., Pirker, K.F., Svergun, D.I., Gautel, M., Djinović-Carugo, K.B., 2014. The structure and regulation of human muscle alpha-actinin. Cell. 159(6), 1447–1460. doi:10.1016/j.cell.2014.10.056.

Rodríguez, S.M., Saslow, L.R., Garcia, N., John, O.P., Keltner, D., 2009. Oxytocin receptor genetic variation relates to empathy and stress reactivity in humans. PNAS. 106(50): 21437–21441. doi:10.1073/pnas.0909579106.

Saevre, C., Meyer, A.M., Van Emon, M.L., Redmer, D.A., Caton, J.S., Kirsch, J.D., Luther, J.S., Schauer, C.S., 2011. Impacts of arginine on ovarian function and reproductive performance at the time of maternal recognition of pregnancy in ewes. Sheep Res Rep. North Dakota State University, Fargo. 52, 13-16. [On line]. Consulting february 23, 2023. Available in: https://pdfs.semanticscholar.org/c540/63c9f5742fd22198d92d5cf367aed3343f01.pdf

Sapolsky, R.M., Romero, L.M., Munck, A.U., 2000. How do glucocorticoids influence stress responses? Integrating permissive, suppressive, stimulatory, and preparative actions. Endocr Rev. 21(1): 55–89. doi:10.1210/er.21.1.55.

Sart, S., Bencini, R., Blache, D., Martin, G.B., 2004. Calm Ewes Produce Milk With More Protein Than Nervous Ewes. Anim Prod Aus. 25, 307.

Selye, H., 1939. The effect of adaptation to various damaging agents in the female sex organs in the rat. Endocr. 25(4), 615–624.

Shen, W., Wang, Q.W., Liu, Y.N., Marchetto, M.C., Linker, S., Lu, S.Y., Chen, Y., Liu, C., Guo, C., Xing, Z., Shif, W., Kelsoeg, J.R., Alda, M., Wang, H., Zhong, Y., Sui, S.F., Zhao, M., Yang, Y., Mi, S., Cao, L., Gage, F.H., Yao, J., 2020. Synaptotagmin-7 is a key factor for bipolar-like behavioral abnormalities in mice. 117(8), 4392.4399. doi:10.1073/pnas.1918165117/-/DCSupplemental.

Sherman, B., Hao, M., Qiu, J., Jiao, X., Baseler, M.W., Clifford Lane, H., Imamichi, T., & Chang, W., 2022. DAVID: a web server for functional enrichment analysis and functional annotation of gene lists. Nucleic Acids Research, v6.8 (Version 6.8). Volume 50, Issue W1, Pages W216–W221, 10.1093/nar/gkac194.

Sigdel, A., Bisinotto, R.S., Peñagaricano, F., 2021. Genes and pathways associated with pregnancy loss in dairy cattle. Sci Rep. 11, 13329. doi:10.1038/s41598-021-92525-0.

Speck, P.A., Collingwood, K.M., Bardsley, R.G., Tucker, G.A., Gilmour, R.S., Buttery, P.J., 1993. Transient changes in growth and in calpain and calpastatin expression in ovine skeletal muscle after short-term dietary inclusion of cimaterol. Biochim. 75(10), 917–923. doi:10.1016/0300-9084(93)90049-x. PMID: 7906151.

Stranden, I., Garrick, D.J., 2009. Technical note: Derivation of equivalent computing algorithms for genomic predictions and reliabilities of animal merit. J. Dairy Sci. 92(6), 2971–2975. doi:10.3168/jds.2008-1929.

Sutherland, M.A., Rogers, A.R., and Verkerk, G.A., 2012. The effect of temperament and responsiveness towards humans on the behavior, physiology and milk production of multi-parous dairy cows in a familiar and novel milking environment. Physiological Behaviour. 107, 329–337. 10.1016/j.physbeh.2012.07.013.

Tanda, G., Ebbs, AL., Kopajtic, T.A., Elias, L.M., Campbell, B.L., Newman, A.H., Katz, J.L., 2007. Effects of muscarinic M-1 receptor blockade on cocaine-induced elevations of brain dopamine levels and locomotor behavior in rats. J Pharmacol Exp Ther. 321(1), 334–344. doi:10.1124/jpet.106.118067.

Tan, P., Allen, J.G., Wilton, S.D., Akkari, P.A., Huxtable, C.R., Laing, N.G., 1997. A splice-site mutation causing ovine McArdle’s disease. Neuromuscul Disord. 7(5), 336–342. doi:10.1016/S0960-8966(97)00062-X.

The UniProt Consortium, UniProt: the universal protein knowledgebase in 2021, Nucleic Acids Research, Volume 49, Issue D1, 8 January 2021, Pages D480–D489. doi:10.1093/nar/gkaa1100

Tost, H., Kolachana, B., Hakimi, S., Lemaitre, H., Verchinski, B.A., Mattay, V.S., Weinberger, D.R., Meyer-Lindenberg, A., 2010. A common allele in the oxytocin receptor gene (OXTR) impacts prosocial temperament and human hypothalamic-limbic structure and function. Proc Natl Acad Sci. 107, 13936– 13941. doi:10.1073/pnas.1003296107.

Van Lier, E., Hart, K.W., Viñoles, C., Paganoni, B., Blache, D., 2017. Calm Merino ewes have a higher ovulation rate and more multiple pregnancies than nervous ewes. Anim. 11(7), 1196–1202. doi:10.1017/S1751731117000106.

VanRaden, P.M., 2008. Efficient methods to compute genomic predictions. J. Dairy Sci. 91(11), 4414–4423. doi:10.3168/jds.2007-0980.

Veissier, I., Boissy. A., Désiré, L., Greiveldinger, L., 2009. Animals emotions: studies in sheep using appraisal theories. Anim Welfare. 18(4), 347–54. doi:10.1017/S0962728600000749.

Voisinet, B.D., Grandin, T., Tatum, J.D., O’Connor, S.F., Struthers, J.J., 1997. Feedlot Cattle with Calm Temperaments Have Higher Average Daily Gains Than Cattle with Excitable Temperaments. J Anim Sci. 75(4), 892– 896. doi: 10.2527/1997.754892x.

Von Borell, E., Dobson, H., Prunier, A., 2007. Stress, behaviour and reproductive performance in female cattle and pigs. Horm Behav. 52(1), 130–138. doi: 10.1016/j.yhbeh.2007.03.014

Walker, K.R., 2006. Characterisation of the Ovine Model of McArdle’s Disease: Development of Therapeutic Strategies. PhD Thesis, Murdoch University.

Wang, H., Misztal, I., Aguilar, I., Legarra, A., Muir, W.M., 2012. Genome-wide association mapping including phenotypes from relatives without genotypes. Genet Res. 94(2), 73–83. doi:10.1017/S0016672312000274.

Wang, H., Misztal, I., Aguilar, I., Legarra, A., Fernando, R.L., Vitezica, Z., Okimoto, R., Wing, T., Hawken, R., Muir, W.M., 2014. Genome-wide association mapping including phenotypes from relatives without genotypes in a single-step (ssGWAS) for 6-week body weight in broiler chickens. Front Genet. 5, 1-10. doi:10.3389/fgene.2014.00134.

Wulf, D.M., Emnett, R.S., Leheska, J.M., Moeller, S.J., 2002. Relationships among glycolytic potential, dark cutting (dark, firm, and dry) beef, and cooked beef palatability. J Anim Sci. 80(7), 1895–1903. doi:10.2527/2002.8071895x.

Wu, G., 2010. Functional amino acids in growth, reproduction, and health. American Society for Nutrition. Adv Nutr. 1(1), 31–37. doi:10.3945/an.110.1008.

Zambra, N., Gimeno, D., Blache, D., Van Lier, E., 2015. Temperament and its heritability in Corriedale and Merino lambs. Anim. 9(3), 373–379. doi:10.1017/S1751731114002833.

